# A Y-specific translocated *gsdf* copy is a candidate master sex-determining gene in common dab (*Limanda limanda*)

**DOI:** 10.64898/2026.11.04.764161

**Authors:** Morten Barvik, Tomasz Furmanek, Prescilla Perrichon, Carl-Johan Rubin, Rolf Brudvik Edvardsen

**Affiliations:** Institute of Marine Research, Bergen, Norway; Institute of Marine Research, Austevoll Research Station, Storebø, Norway; Department of Medical Biochemistry and Microbiology; Uppsala University, Uppsala, Sweden

## Abstract

Sex-determining systems in flatfishes are highly diverse, and even closely related species can differ in both sex-determining loci and heterogametic systems. In common dab (*Limanda limanda*), the genetic basis of sex determination has not yet been established. Here, we investigated sex determination using pooled whole-genome sequencing combined with long-read genome sequencing of females and males from Iceland and Norway. Genome-wide allele-frequency analyses identified sex-associated regions consistent with male heterogamety (XX/XY), with the strongest signal on chromosome 16. Comparison of the existing reference assembly, which represents an X haplotype at this locus, with a haplotype-resolved male PacBio HiFi assembly revealed a ∼21.5 kb Y-specific insertion on chromosome 16. The insertion showed complete coverage in male pools but little or no coverage in female pools from either population. It contained a translocated genomic segment including a copy of gonadal soma-derived factor (*gsdf*) and a predicted gene of unknown function. Comparative genomic analyses were consistent with this segment originating from an autosomal *gsdf*-containing region on chromosome 1. Given the recurrent recruitment of *gsdf* as a sex-determining factor in teleosts, we designate the Y-linked copy *gsdfY* and identify it as the strongest candidate for the master sex-determining gene in common dab. Additional broad sex-associated differentiation on chromosome 19 was restricted to the Icelandic samples and may reflect population-specific expansion of the sex-linked region, although its structural basis remains unresolved. Together, these findings provide the first genomic characterization of sex determination in common dab and demonstrate how gene duplication and translocation can contribute to the evolution of novel sex-determining systems in flatfishes.

## Introduction

Vertebrates have evolved different mechanisms for determining sex, ranging from strictly genetic sex-determining systems to systems where environmental influences determine sex [1]. In genetic sex-determining systems, sex is primarily determined by a master sex-determining (MSD) gene or locus that directs the gonadal development towards the formation of either ovaries or testes [2]. Comparatively, other sex-determining systems may be strongly influenced by environmental factors, and thus genetic differentiation between sexes can be weak or absent [3]. Unlike mammals and birds, teleost fishes display remarkable plasticity in sex determination, including frequent turnover of sex chromosomes, recruitment of novel MSD genes and sensitivity to environmental cues such as temperature [4–6]. This rapid turnover often results in evolutionarily young sex chromosomes with low genetic differentiation between X and Y or Z and W haplotypes [7,8]. Despite the diversity of sex-determining systems in teleosts, the same developmental pathways are repeatedly recruited when determining sex. In many fish species, MSD genes are derived from genes with already conserved roles in gonadal development, particularly genes involved in testis differentiation. Genes influencing the transforming growth factor beta (TGF-β) signalling pathway are often identified as the MSD gene in fishes [9–11]. For example, *gsdf* has been identified as the MSD gene in Atlantic halibut [12], *amh* has been reported as the MSD gene in Japanese flounder/olive flounder [13], and *bmpr1ba* has been suggested as a putative MSD gene in Pacific halibut [14]. These examples show that novel sex-determining systems in fishes can evolve through recruitment of genes that already have an established function in the gonadal differentiation pathways. New sex-determining loci can arise through several mechanisms, including allelic change, change in regulatory environment, gene duplications and larger structural rearrangements. Duplications or translocations in the genome involving gonadal development genes, can lead to new genomic environments, which might in turn alter gene dosage or place a gene under new regulatory control. These processes can provide genes that already have an established function in the downstream gonadal differentiation pathways with a new role as an upstream master sex-determining factor.

The flatfishes represent an order where this reproductive plasticity is readily apparent. Genomic and experimental studies have shown that closely related flatfish species can differ in sex-determining mechanisms, including female heterogametic ZZ/ZW systems, male heterogametic XX/XY systems and systems with strong environmental influences [8,15–17]. Within flatfishes, members of the family Pleuronectidae provide examples of *gsdf*, *amh* and *bmpr1ba* as master or candidate sex-determining genes in different flatfish lineages, indicating that sex determination in this family has not followed a conserved evolutionary trajectory. This diversity suggests that pleuronectid flatfishes have experienced extensive turnover in sex-determining systems during their evolutionary history.

The common dab *(Limanda limanda)* is a pleuronectid flatfish native to coastal waters in the northeastern Atlantic Ocean. The genetic basis of sex determination has not been described in common dab. Given the diversity of sex-determining systems among related flatfishes, investigating this species may provide further insight into the evolution of sex determination in flatfishes. In this study we used whole-genome pool-sequencing and a male long-read genome assembly to investigate genomic differences between female and male common dab and identify candidate sex-determining loci. Our results provide insight towards understanding the sex-determining mechanism of this species and establishes *gsdf* as a potential candidate MSD gene.

## Materials and Methods

### Ethics Statement

All procedures involving live animals were approved by the Norwegian Food Safety Authority (FOTS number: 30644). Handling of live animals, rearing of wild-caught fish, and euthanasia before sampling followed the principles of the Norwegian Animal Welfare Act.

### Sample collection and establishment of phenotypic sex

A total of 40 common dab individuals were collected during a research survey in the coastal region of northwestern Iceland. Individuals were euthanized by forcefully driving a sharp blade through their brain. Phenotypic sex was assigned by dissection and examination of the gonad based on established morphological criteria. Whole pectoral fins, or sections of pectoral fins, were excised shortly after euthanasia. Fin clips from 20 females and 20 males were collected and stored in absolute ethanol.

In addition, 18 wild common dab individuals were collected around the archipelago of Austevoll, Norway, then transported to the Austevoll research station, Institute of Marine Research, Storebø, Norway. Individuals were then euthanized with a lethal dose of tricaine methanesulfonate (MS-222). Phenotypic sex was assigned by gonadal examination, and 9 females and 9 males were sampled. Gill samples from these individuals were excised, flash-frozen in liquid nitrogen and stored at -80 °C. One male individual was selected for long read sequencing. Heart tissue was excised from this individual, flash-frozen in liquid nitrogen and stored at -80 °C.

### DNA extraction and whole genome sequencing

Genomic DNA from the 40 Icelandic individuals was extracted from fin clips using the DNAdvance extraction reagent kit (Beckman Coulter, Inc., Brea, CA, USA) on a Biomek i5 automated pipetting workstation (Beckman Coulter Life Sciences, Indianapolis, IN, USA), following the manufacturer’s instructions. DNA quantity and quality were assessed using Nanodrop 8000 Spectrophotometer (Thermo Fisher Scientific, Wilmington, DE, USA) and Qubit Flex Fluorometer (Invitrogen, Thermo Fisher Scientific, Singapore) measurements. Extracted DNA from fin clips of Icelandic individuals was divided into female and male pools containing 10 individuals each, producing two Icelandic female pools and two Icelandic male pools.

Genomic DNA from the 18 Norwegian individuals was manually extracted from gill tissue using the DNeasy Blood & Tissue Kit (QIAGEN, Hilden, Germany), following the manufacturer’s instructions. DNA quantity and quality were assessed using NanoDrop and Qubit measurements. Extracted DNA from Norwegian individuals was divided into two pools, one pool of 9 females and one pool of 9 males.

All 6 pools (Icelandic fin clip pools and Norwegian gill pools) were sequenced on one NovaSeq 6000 SP flow cell (Illumina, Inc., San Diego, CA, USA), with 2 x 150 bp paired-end sequencing. Libraries were prepared according to the TruSeq DNA Nano Library Prep protocol (Illumina, Inc., San Diego, CA, USA). Reads were adapter trimmed and aligned to the common dab reference genome (GCF_963576545.1). Mean mapped sequencing depth ranged from approximately 60x to 70x across the six pools.

### Male genome assembly

To supplement the common dab reference genome, heart tissue from a Norwegian male individual was used to generate a male genome assembly. Tissue from this individual was sent to Novogene UK Company Ltd (Cambridge, UK) for high-molecular-weight genomic DNA extraction and PacBio HiFi sequencing. DNA extraction, library preparation, and sequencing were performed by Novogene according to their standard protocols. Raw HiFi reads were assembled using hifiasm version 0.25.0-r726 with default settings [18]. The final diploid assembly contig sets had lengths of 719 and 671 Mb, respectively for haplotype 1 and haplotype 2 contigs. Hap1 consisted of 418 contigs while hap2 consisted of 224 contigs with contig N50 values of 19.6 and 20.9 Mb, respectively. GC content was 42.6% for haplotype 1 and 42.4% for haplotype 2. Assembly statistics, including total assembly size, number of contigs, contig N50 and GC content, were calculated using QUAST [19]. Sequence alignments were performed by minimap2 version v2.30 with default settings [20].

### Identification of sex-associated SNPs

Sequence variation between phenotypic females and males was assessed by comparing pooled female samples from Norway and Iceland with pooled male samples from Norway and Iceland. Pools were sequenced on an Illumina sequencer to generate paired-end reads. Resulting reads were adapter trimmed and aligned to both the common dab reference genome and haplotype 2 of the male PacBio HiFi assembly generated in this study. Both haplotypes were initially examined. Haplotype 2 was used for the subsequent analyses because it contained the male-specific chromosome 16 insertion and therefore represented the putative Y haplotype at this locus. Read mapping was performed with bwa-mem2 [21] using default paired-end alignment parameters. SAM files were converted to BAM format, coordinate sorted and indexed using SAMtools version 1.22 [22]. Variant detection was performed separately for the common dab reference genome and the male PacBio HiFi assembly. The allele count for each genomic position was extracted from the aligned BAM files using SAMtools mpileup. Pileup generation was restricted to high confidence bases and alignments by applying a minimum base- and mapping-quality score of 20. Sites with low read depth were excluded. Only biallelic SNPs were retained for downstream analysis. For each retained SNP, reference and alternative allele counts were counted separately for each female and male pool. Minor allele frequencies were calculated as the proportion of reads supporting the less frequent allele relative to the total read depth at each genomic position. To identify sex-associated SNPs consistent with a male heterogametic XY system, the retained SNPs from the pileup were filtered for loci that were fixed in female pools but had a varying degree of allele frequency in male pools. Loci fixed in females were required to have a minor allele frequency below 0.01 in each female pool. Retained SNPs were also filtered to have a read depth of 20 reads or higher to be included. All the SNPs that passed these filters were eligible to be considered as candidate sex-associated markers if the corresponding loci in the male pools showed a high minor allele frequency, with the strongest candidates expected to approach a male minor allele frequency of 0.5. For each candidate SNP the male minor allele frequency was calculated per pool and averaged across male pools as the mean male minor allele frequency.

Resulting candidate sex-associated SNPs were visualized as Manhattan style plots across the common dab reference genome and the male PacBio HiFi assembly. Genomic coordinates were plotted according to their position in the reference genome or the male haplotype 2, and the calculated mean male minor allele frequency was used to differentiate between sex-linked SNPs. Regions with a high abundance of SNPs that showed an elevated mean male minor allele frequency were established as candidate sex-associated genomic regions.

### Data availability

All data supporting the results of this study are included in the article and its supplementary information. The sequencing data has been uploaded to the European Nucleotide Archive under project PRJEB127433.

## Results

### Pool-seq identifies sex-linked markers compatible with an XX/XY system in common dab

Whole-genome short-read sequencing was performed on 58 phenotypically sexed common dab, comprising 40 individuals from Iceland and 18 from Norway. The Icelandic samples were divided into two female and two male pools of 10 individuals each, while the Norwegian samples were divided into one female and one male pool of nine individuals each. Reads from all six pools were aligned to the common dab reference genome, and genome-wide SNP differentiation between female and male pools was assessed to identify sex-associated genomic regions. The sex of the individual used to generate the reference genome is recorded as “not collected” in the associated NCBI BioSample. However, the Y-specific chromosome 16 insertion identified here is absent from both the primary assembly and the alternate haplotype, suggesting that the reference individual was female.

Under male heterogamety, Y-linked sequence variants are expected to occur as male-specific alleles, resulting in minor allele frequencies approaching 0.5 in male pools while remaining absent or near zero in female pools. Minor allele frequency was calculated as the frequency of the less abundant allele at each biallelic site and therefore ranged from 0 to 0.5. SNPs displaying allele-frequency patterns compatible with either male (XX/XY) or female (ZZ/ZW) heterogamety were identified and evaluated. A total of 3,985,469 SNPs passed filtering criteria and displayed SNPs with a male minor allele frequency > 0 and ≤ 0.5. Female heterogamety was evaluated using the reciprocal criterion. Here 89,828 SNPs passed the filtering criteria.

When female heterozygous SNPs are plotted on a genome-wide Manhattan plot, we saw no enriched regions for a potential ZZ/ZW system. When male heterozygous SNPs are plotted, three regions enriched for sex-associated SNPs were detected across the common dab reference genome, located on chromosomes 1, 16, and 19 (Fig 1A). Among these, the strongest signal was observed on chromosome 16, where a pronounced peak of male-specific differentiation spans approximately 12.4 Mb to 13.8 Mb (Fig 1C). In addition to the major peak, chromosome 16 contained a broader region enriched for sex-associated SNPs, extending from approximately 14 Mb to the end of the chromosome.

**Fig 1.**
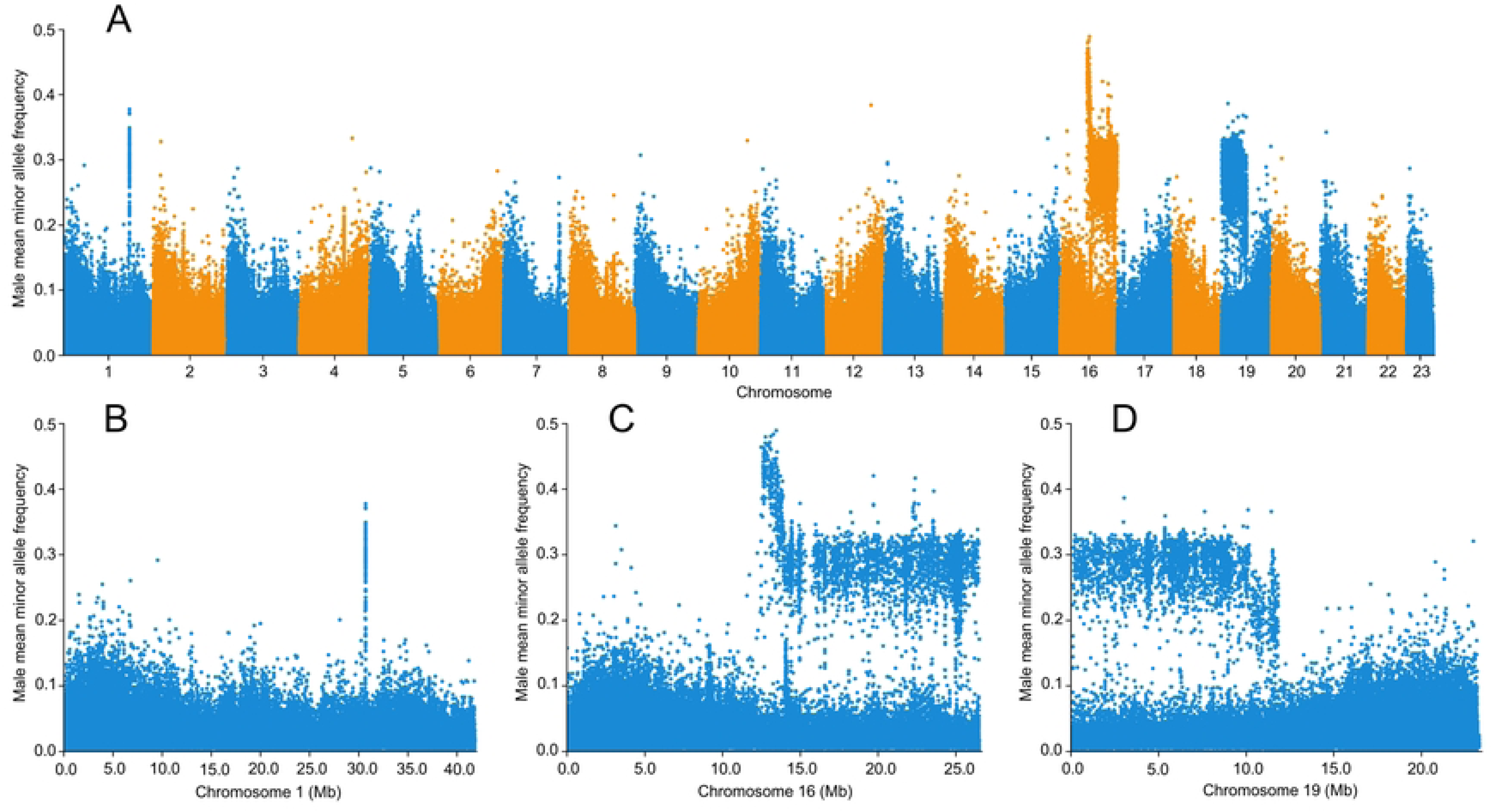
Genome-wide identification of sex-associated SNPs in common dab using the common dab reference genome. (A) SNPs with an XX/XY compatible allele frequency pattern across the common dab reference genome assembly (GCF_963576545.1). The y-axis shows male mean minor allele frequency across male pools from both Iceland and Norway. The strongest sex-associated signal is located on chromosome 16, with additional signals on chromosomes 1 and 19. (B) Detailed view of chromosome 1 showing a narrow peak of sex-associated SNPs near 30 Mb. (C) Detailed view of chromosome 16 showing a pronounced peak of male-specific differentiation between approximately 12.4 and 13.8 Mb, together with a broader region enriched for sex-associated SNPs extending towards the end of the chromosome. (D) Detailed view of chromosome 19 showing an extended region of sex-associated SNPs spanning from the start of the chromosome to approximately 12 Mb.

A second broad signal was detected on chromosome 19 (Fig 1D). This signal resembles the broad differentiated region observed on chromosome 16. On chromosome 19 we observe differentiated SNPs distributed across a broad region spanning almost 12 Mb starting at one end of the chromosome.

A third signal was detected on chromosome 1 (Fig 1B) near 30 Mb. This narrow signal of differentiated SNPs contrasts the wide differentiated regions observed on chromosomes 16 and 19.

### Male long read sequencing confirms chromosomes 16 and 19 as potential sex chromosomes

To complement the available common dab reference genome and enable direct characterization of male genomic sequences, a male individual was sequenced using PacBio HiFi long-read sequencing and a haplotype-resolved male assembly was generated. To validate the sex-associated signals identified against the common dab reference and to investigate the male haplotype in greater detail, the allele frequency analysis was repeated using haplotype 2 of the male PacBio HiFi assembly (Fig 2). Haplotype 2, and not haplotype 1, contained the sex-associated insertion described below and was therefore used for the male-assembly-based pool-seq analysis. The same pooled libraries of female and male samples were aligned to the male haplotype 2 and SNP differentiation between female and male pools were assessed at a genome-wide level. The same filtering parameters were used as in the common dab reference analysis. The analysis identified 3,900,580 SNPs displaying an XX/XY compatible allele frequency pattern.

**Fig 2.**
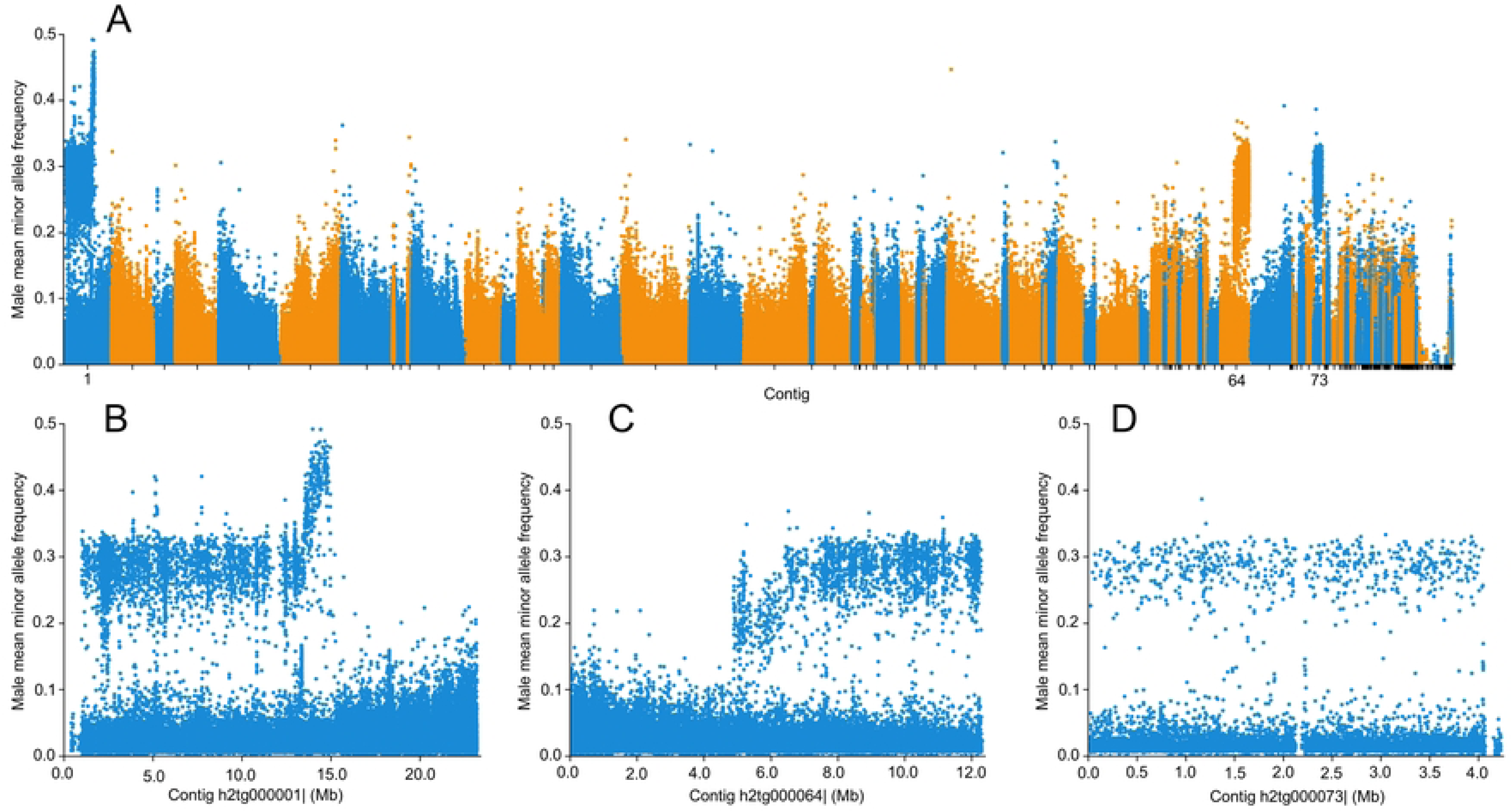
Genome-wide identification of sex-associated SNPs in common dab using haplotype 2 of the male genome assembly. (A) SNPs with an XX/XY compatible allele frequency pattern across haplotype 2 of the male common dab assembly generated in this study. The strongest sex-associated signal is located on contig 1 (h2tg000001), which is syntenic to chromosome 16 in the common dab reference assembly. Additional signals are present on contigs 64 and 73 (h2tg000064 and h2tg000073), corresponding to chromosome 19 in the common dab reference assembly. (B) Detailed view of contig 1 showing a major peak of sex-associated SNPs between approximately 13.5 and 14.8 Mb, together with a broader region enriched with sex-associated SNPs spanning from almost the start of the contig to the major sex-associated peak. (C) Detailed view of contig 64 showing a region of sex-associated SNPs from approximately 5 Mb, extending out to the end of the contig. (D) Detailed view of contig 73 showing elevated male-specific differentiation across the entire contig.

The analysis of the male haplotype 2 provided results consistent with the analysis of SNPs across the common dab reference genome (Fig 2A). The most prominent signal was located on contig 1, which aligns to chromosome 16 in the common dab reference genome (S1 Fig). Contig 1 contains a pronounced peak of male-specific differentiation from approximately 13.5 to 14.8 Mb (Fig 2B), corresponding to the signal identified on chromosome 16 in the common dab reference analysis. In addition to this localized peak, contig 1 exhibits a broader region enriched with sex-associated SNPs as seen on chromosome 16 in the common dab reference genome. This consensus between the male haplotype 2 and the common dab reference genome supports that chromosome 16 exhibits strong genomic differentiation between females and males.

Signals corresponding to chromosome 19 in the common dab reference genome were also identified in the male haplotype 2. In the male haplotype 2 these signals are distributed across contigs 64 and 73, which together correspond to chromosome 19 in the common dab reference genome (Fig 2C-D, S1 Fig). On contig 64, sex-associated SNPs appear from approximately 5 Mb and continue to the end of the 12 Mb contig. On contig 73 there are sex-associated SNPs covering almost the entire length of the 4 Mb contig. The sex-associated patterns observed on contigs 64 and 73 correspond to the chromosome 19 signal in the common dab reference genome, supporting the presence of sex-linked differentiation on chromosome 19.

The narrow signal observed on chromosome 1 in the common dab reference genome was not recovered in the analysis of the male haplotype 2, this is most likely explained by reads from the Y-specific *gsdf* duplication mapping to the homologous *gsdf* region on chromosome 1 as described below. Overall, the analysis of the male haplotype 2 confirmed that there exists sex-linked differentiation between females and males on chromosomes 16 and 19.

### Male-specific insertion within the sex-associated region on chromosome 16

The strongest sex-associated signal identified by the analysis of both the common dab reference and the male haplotype 2 is the region on chromosome 16 spanning from 12.4 Mb to 13.8 Mb. To investigate whether the signal was associated with structural variation between females and males, read-depth patterns in the corresponding region of the male haplotype 2 were compared between the female and male pools. This revealed an approximately 21.5 kb sequence present in the male haplotype 2, which is not found in the common dab reference genome. This male-specific inserted sequence appears at ∼14.8 Mb on contig 1 on the male haplotype 2, within the corresponding signal found on chromosome 16 (Fig 3).

**Fig 3.**
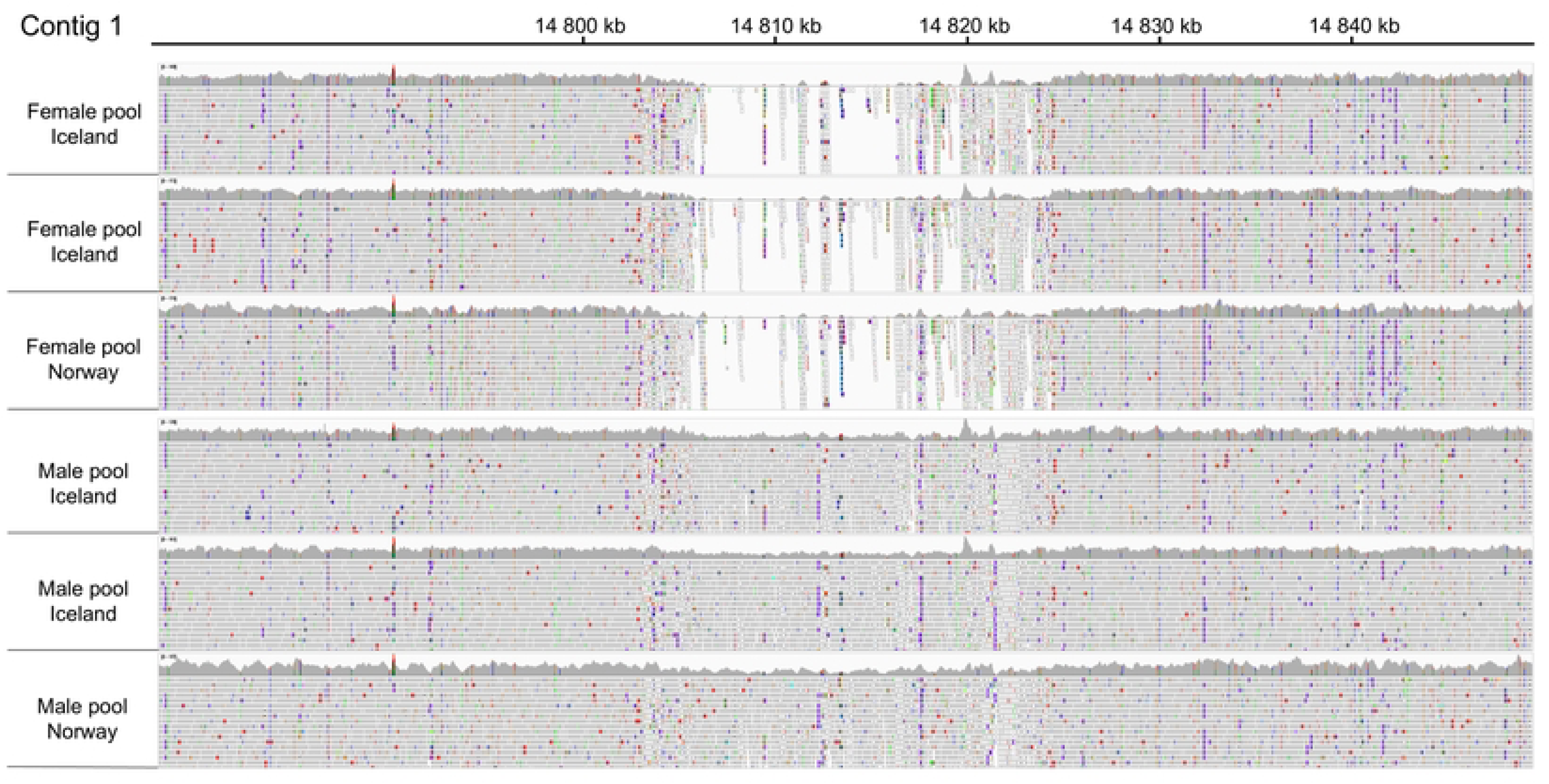
Male-specific insertion within the sex-associated region on contig 1. Read coverage across a male-specific insertion on contig 1 at approximately 14.8 Mb, visualized in IGV. The upper tracks show female pools from Iceland and Norway, and the lower tracks show males from Iceland and Norway. Male pools show continuous read coverage across an approximately 21.5 kb long inserted sequence and its flanking regions, while female pools show little to no coverage across the insertion.

The inserted sequence showed a clear sex-specific pattern. Reads from the male pools mapped continuously across the inserted region and its flanking sequences, and reads from female pools showed little to no coverage across the inserted region. This pattern was observed across female and male pools from both Iceland and Norway, indicating that this male-specific insertion is present in different geographical populations. The presence of male reads spanning this region, and the corresponding absence of female reads, is consistent with a XX/XY sex-determining system where this inserted sequence is located on a male-specific Y haplotype and is absent from the corresponding X haplotypes.

The location of this inserted sequence in the male haplotype 2 corresponding to the strongest sex-associated signal found on chromosome 16 suggest that this 21.5 kb inserted sequence is a strong candidate for a Y-specific structural variant in common dab.

### The male-specific insertion contains a duplicated *gsdf* copy

Inspection of the common dab genome annotation identified three intact autosomal *gsdf*-like genes. Two tandemly arranged copies on chromosome 1 are annotated as growth/differentiation factor 6-like (NCBI Gene IDs 133003044 and 133004413), whereas a third copy on chromosome 5 is annotated as bone morphogenetic protein 6-like (Gene ID: 133002334). Because automated annotations of teleost *gsdf* genes are inconsistent, and because these genes share high sequence similarity and the same five-exon structure, we hereafter refer to them as *gsdf1*, *gsdf2* and *gsdf3*, respectively (Fig 4A). Comparative synteny analysis showed that *gsdf1* and *gsdf2* are located within a conserved genomic region containing *scarb2*, *nup54*, *ppef2*, *gsdf*, *aff1* and *mrc1* (S2 Fig). Tandemly arranged *gsdf* copies were also present at this locus in several other flatfish species, consistent with the chromosome 1 region represents the ancestral *gsdf* locus and that the tandem duplication predates the divergence of common dab from several related flatfish lineages.

**Fig 4.**
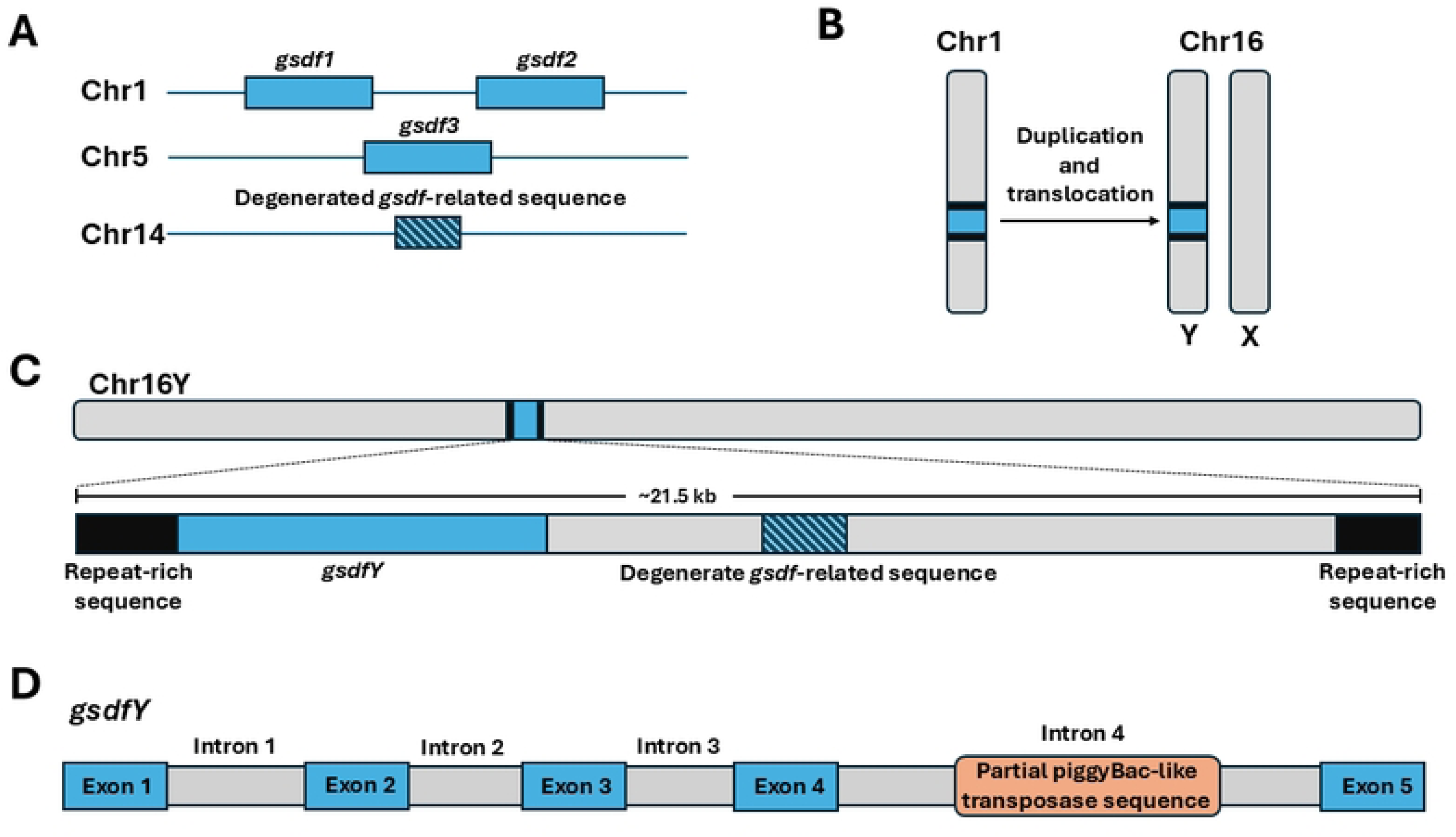
Origin and structure of *gsdfY* in the male-specific insertion on contig 1. Only *gsdf*-related features and flanking repeat-rich sequences are shown; the conserved *LOC132999031-like* gene within the insertion is omitted. The schematics in this figure are not drawn to scale. (A) Genomic distribution of *gsdf*-related sequences identified in common dab. Two copies, designated *gsdf1* and *gsdf2*, occur as a tandem duplication on chromosome 1. An additional copy is located on chromosome 5, designated *gsdf3*, and one degenerate *gsdf*-related sequence is found on chromosome 14. (B) Schematic representation of the proposed duplication and translocation event. A segment on chromosome 1 containing the tandem *gsdf* region has been duplicated and translocated into the chromosome 16 male-specific Y haplotype. (C) Structure of the approximately 21.5 kb male-specific insertion on chromosome 16. The inserted segment contains one apparently complete *gsdf* copy most closely related to *gsdf1,* designated *gsdfY*, together with a degenerated *gsdf*-related sequence. (D) Structure of the *gsdfY* gene. BLAST analysis identified a sequence fragment within the expanded intron showing high similarity to piggyBac-like transposable elements.

The sequence content and the genomic origin of the ∼21.5 kb insertion was examined to pinpoint potential candidate sex-determining sequences within the insert. Sequence alignment identifies that the male-specific insertion on chromosome 16 corresponds to a duplicated and translocated region originating from the *gsdf*-containing region on chromosome 1 (Fig 4B). Because the male assembly was haplotype-resolved, both haplotype assemblies were inspected. The sex-associated insertion containing the duplicated *gsdf* copy was located only on haplotype 2.

This apparently intact copy of *gsdf* inside the male-specific inserted region represents the strongest candidate sex-determining gene identified in this study and will be referred to as *gsdfY*. The chromosome 16 insertion also contains a predicted *LOC132999031-like* gene encoding an uncharacterized protein. Two related copies occur near *gsdf1* and *gsdf2* on chromosome 1, one near *gsdf3* on chromosome 5, and a partial copy near the degenerate *gsdf*-related sequence on chromosome 14. The conserved association between these genes suggests that the chromosome 16 insertion originated through duplication and translocation of a larger genomic segment rather than of *gsdf* alone.

Alignment of the complete coding sequences showed that *gsdfY* was most similar to *gsdf1*, differing at only three nucleotide positions and showing 99.5% nucleotide and 98.5% predicted protein sequence identity. By comparison, *gsdfY* showed 98.0% nucleotide and 96.5% protein sequence identity to *gsdf2*, and 96.2% nucleotide and 94.5% protein sequence identity to *gsdf3* (S3 Fig and S1 Table). All four intact *gsdf* copies shared a five-exon structure. Intron 4 was 778, 464 and 606 bp long in *gsdf1*, *gsdf2* and *gsdf3*, respectively, but was expanded to approximately 1685 kb in *gsdfY*. BLAST analysis identified a sequence fragment within the expanded intron showing high similarity to piggyBac-like transposable elements. Related sequences were present at several other positions in the common dab genome but were absent from the corresponding introns of the autosomal *gsdf* copies.

### Icelandic samples show sex-associated differentiation on chromosomes 16 and 19

Genome-wide analysis of sex differentiated SNPs exclusively from the pools originating from Iceland displays two distinct clusters of sex-linked SNPs on two different chromosomes, 16 and 19 (Fig 5A). This analysis was performed on data from pools originating from Iceland, excluding pools from Norway. Sequenced data from the Icelandic pools were mapped to the common dab reference genome. The two regions of high SNP differentiation on chromosomes 16 and 19 correspond to the signals identified in the combined analysis of all pools (Figs 1A and 2A). When the analysis is restricted to the Icelandic pools, these signals become more pronounced (Fig 5B–C).

**Fig 5.**
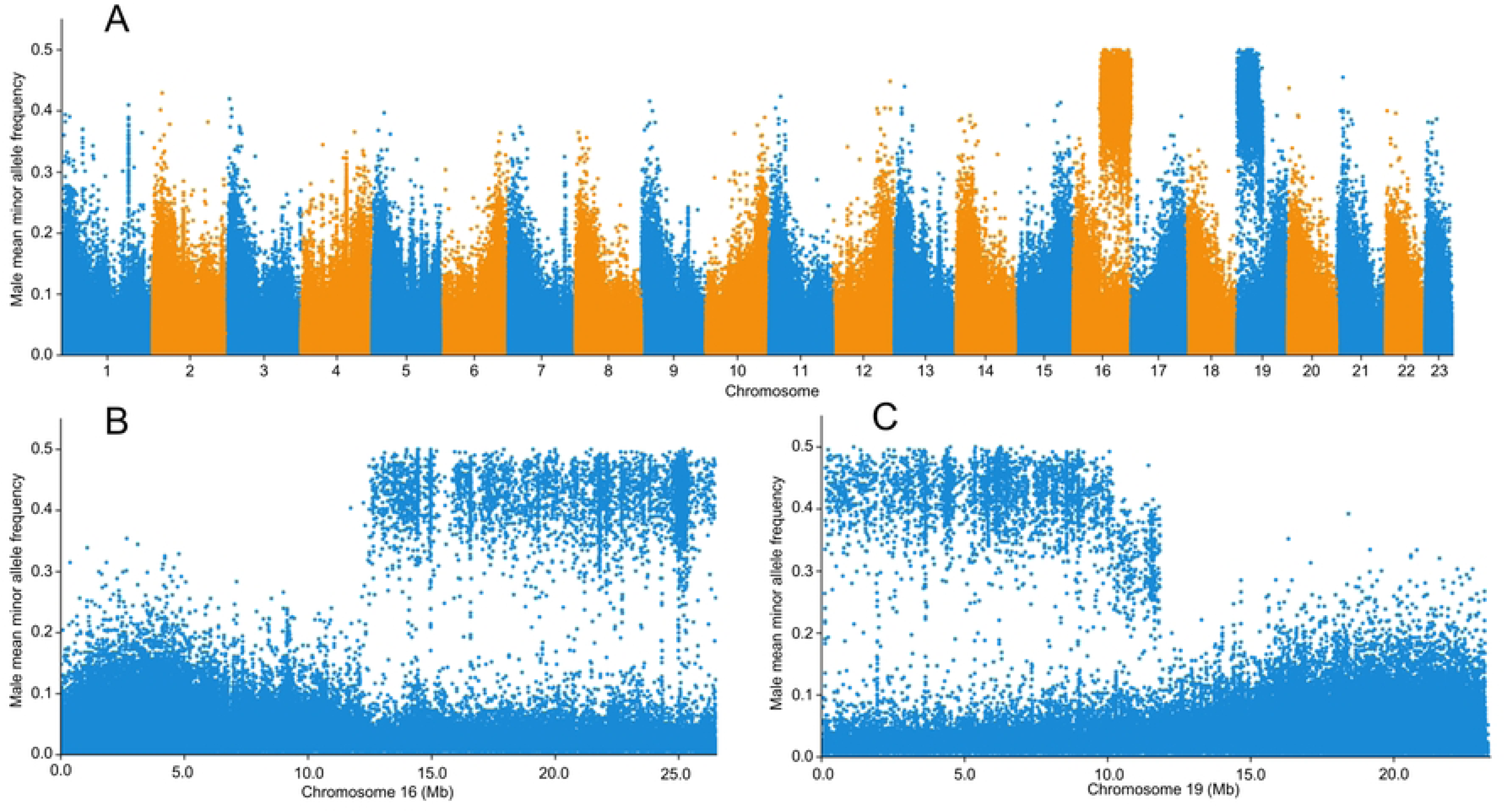
Extended sex-associated differentiation on chromosomes 16 and 19 in Icelandic common dab pools. (A) SNPs with an XX/XY compatible allele frequency pattern across the common dab reference genome assembly (GCF_963576545.1). The y-axis shows male mean minor allele frequency across male pools from Iceland. Two major regions of sex-associated SNPs are detected on chromosomes 16 and 19. (B) Detailed view of chromosome 16 showing extended sex-associated differentiation including the primary chromosome 16 sex-associated interval identified with pools from both Iceland and Norway. (C) Detailed view of chromosome 19 showing a broad and continuous region of sex-associated SNPs spanning approximately half the length of the chromosome.

## Discussion

Our results support a male-heterogametic XX/XY sex-determining system in common dab and identify a Y-specific *gsdf* copy, generated by duplication and translocation, as the putative master sex-determining gene. Genome-wide comparisons between female and male pools revealed a sex-associated region on chromosome 16. Within this region, the male haplotype 2 contained an approximately 21.5 kb insertion that was covered completely by reads from males but showed little or no coverage in females from either Iceland or Norway. The insertion contains an apparently intact copy of *gsdf*, here designated *gsdfY*, together with a degenerated *gsdf*-related sequence. Sequence comparisons across the common dab reference genome suggest that the male-specific insertion originated from chromosome 1. Chromosome 1 contains a region with an autosomal tandem *gsdf*-duplication comprising *gsdf1* and *gsdf2*. Synteny between selected flatfish species reveals that this tandem *gsdf*-duplication is present in some, but not all flounders (S2 Fig). This tandem duplication is not sex-specific and therefore not sex determining. By contrast, the insertion on chromosome 16 is restricted to the Y haplotype and contains a copy most closely related to *gsdf1*, together with a degenerated *gsdf* sequence. The most parsimonious interpretation is therefore that the tandem duplicated *gsdf*-sequences on chromosome 1 were copied and translocated to chromosome 16, where the *gsdf1*-derived copy became the Y-linked candidate sex-determining gene, *gsdfY*. The conserved syntenic position of the chromosome 1 tandem copies indicates that this region represents the ancestral *gsdf* locus in common dab. The high sequence similarity between *gsdfY* and *gsdf1* further suggests that *gsdfY* originated from this locus, although the precise donor copy cannot be established from coding-sequence similarity alone. The Y-linked copy retained the same five-exon structure but acquired a substantially expanded fourth intron containing a well-conserved piggyBac-like transposable-element fragment. This raises the possibility that transposable-element activity contributed to the duplication, translocation or subsequent structural evolution of *gsdfY*. However, because the fragment is incomplete and located within the gene rather than at the boundaries of the chromosome 16 insertion, its mechanistic involvement remains uncertain. The sex-specific occurrence of this structural variation, its location within the strongest sex-associated interval, and the established role of *gsdf* in testis development collectively make *gsdfY* a strong candidate for a master sex-determining gene in common dab. Although the insertion also contains a conserved predicted gene of unknown function, *gsdfY* remains the strongest candidate because of the established role of *gsdf* in testis differentiation and its recurrent recruitment as a master sex-determining gene in teleosts. The identification of *gsdf* as a candidate master sex-determining gene in common dab is not surprising as *gsdf* is actively involved in testis differentiation [23–25] and has acquired a master sex-determining function in several teleost species [26,27]. In Atlantic halibut *gsdf* is the master sex-determining gene, and a genomic region upstream of *gsdf* has been proposed as a regulatory element on the Y chromosome in males [12]. *Gsdf* has acquired sex-determining function in various ways, either through allelic mutation, copy number variation or sex specific cis regulation of *gsdf*. These examples give us insight into how *gsdf* might go from a conserved gene associated with the differentiation of sex to becoming the main switch that determines sex on a higher level.

The narrow chromosome 1 signal was not recovered when reads were mapped to the male haplotype 2 and likely reflects cross-mapping of reads from the Y-specific *gsdf* insertion to the homologous autosomal *gsdf* region in the common dab reference genome.

The details around the sex-determining function of the new *gsdf* duplication remain to be clarified. One possible explanation is that an additional copy of *gsdf* provides a higher *gsdf* expression, resulting in an increased *gsdf* dosage in males. The insertion into chromosome 16 may also have placed *gsdfY* in a novel regulatory environment. This change in expression could mean earlier gene expression, stronger gene expression or changing which tissue the gene is expressed in during a critical window of sex determination in common dab. The retained flanking sequences within the insertion may also contain regulatory elements copied from chromosome 1, whereas the chromosome 16 insertion site may contribute additional cis-regulatory activity. All these alternatives are not mutually exclusive. Previous work in Atlantic halibut has emphasized the effect on sex differentiated regulation of the *gsdf* gene [12] where a Gypsy-like LTR remnant immediately upstream of *gsdf* was proposed to act as a promoter driving early male-specific *gsdf* expression. Developmental expression analyses are needed to determine whether *gsdfY* is expressed before other components of the male differentiation pathway and differs from the autosomal *gsdf* copies in timing, tissue specificity or expression level. Functional validation through targeted knockout in genetic males or knock-in in genetic females would test its proposed role in sex determination.

In addition to the genomic interval on chromosome 16 shared between Icelandic and Norwegian samples, we observe a striking pattern of extensive sexually differentiated SNPs spanning several Mb on chromosomes 16 and 19. A pronounced chromosome 19 signal was detected in the Icelandic pools, whereas no comparable signal was detected in the Norwegian pools under the present analysis. In contrast, the chromosome 16 insertion showed a consistent sex-specific pattern in samples from both geographic regions. These observations suggest that the Y-specific insertion is present within the sampled material from both locations, while the extensive chromosome 16 and chromosome 19 differentiation is primarily associated with the Icelandic samples. The broad sex-associated regions observed only in the Icelandic samples may reflect population-specific sex-chromosome variation rather than components essential to sex determination in common dab. One possible explanation is a fusion between the *gsdfY*-bearing chromosome 16 Y haplotype and chromosome 19. Such a Y–autosome fusion could reduce recombination and extend sex linkage across both chromosomes, generating a larger, population-specific Y chromosome. A putative Y–autosome fusion has similarly been proposed in Greenland halibut, providing a relevant comparison from another pleuronectid flatfish [28]. Although the distribution of the sex-associated regions may be consistent with a fusion, the present data do not demonstrate a physical connection between chromosomes 16 and 19.

Common dab has previously been reported to have a diploid chromosome number of 2n = 46 (23 chromosome pairs) [29]. Di and Knowles did however observe metaphase cells with chromosome numbers below 46, which they attributed to chromosome loss during preparation of the samples. Our observation of sex-associated differentiation involving chromosomes 16 and 19 raises the possibility that structural chromosome polymorphism may occur within common dab, making these earlier observations noteworthy in retrospect and emphasize the value of renewed cytogenetic analysis of common dab.

If confirmed, such a fusion could represent a secondary stage in the evolution of the common dab sex chromosome. In this scenario, the insertion of *gsdfY* into chromosome 16 established an ancestral Y chromosome shared by the Icelandic and Norwegian populations, whereas a subsequent fusion in the Icelandic lineage expanded the sex-linked region to include chromosome 19. This would explain why the localized chromosome 16 signal is shared between populations, while the broader chromosome 19 signal is restricted to Iceland. The limited differentiation surrounding *gsdfY* is also consistent with a relatively young sex-determining region in which X and Y remain largely homologous. Chromosome-scale phased assemblies supported by Hi-C or karyotyping, combined with broader population sampling, will be required to test this model and determine the geographic distribution and evolutionary history of the proposed rearrangement.

In conclusion, our findings identify a male-specific insertion on chromosome 16 containing *gsdfY* as the strongest candidate master sex-determining gene in common dab. The available evidence is consistent with duplication of a pre-existing autosomal *gsdf* region from chromosome 1, followed by translocation to the chromosome 16 Y haplotype. The distinct genomic patterns observed in Icelandic and Norwegian samples further suggest that the extent of sex-associated differentiation may vary among populations, although the structural basis of this variation remains to be determined. Together, these results illustrate how gene duplication, translocation and potentially subsequent chromosome rearrangements can contribute to the evolutionary diversification of sex-determining systems. They also reinforce the view that closely related flatfish lineages have repeatedly recruited different genes, chromosomes and structural variants to establish novel mechanisms of sex determination.

## Acknowledgements

We would like to thank the Marine and Freshwater Institute at Hafnarfjörður, Iceland for giving us the opportunity to guest the research vessel Árni Friðriksson on one of its surveys to collect flatfish samples. We thank Oddmund Skår and Velimir Nola for providing essential support for capturing common dab adults and the technical staff of the Institute of Marine Research, Austevoll Research Station for their assistance in maintaining the fish until sampling.

## Supporting information

**S1 Fig. Synteny between sex-associated male haplotype 2 contigs and the common dab reference genome.** Dot plot showing whole-contig alignments between male haplotype 2 contig 1 and chromosome 16 of the common dab reference assembly (highlighted in blue), and between male haplotype 2 contigs 64 and 73 and chromosome 19 (highlighted in red). The alignments support the assignment of the sex-associated male haplotype 2 contigs to chromosomes 16 and 19 in the reference genome.

**S2 Fig. Synteny of chromosomal regions associated with *gsdf* across selected flatfish species.** Synteny of chromosomal regions associated with *gsdf.* Gene order and orientation is shown to illustrate conserved features of the *gsdf* genomic neighbourhood. Solid green arrows represent *gsdf* genes, whereas hatched arrows represent *LOC132999031-like* genes encoding predicted proteins of unknown function. Tandemly duplicated *gsdf*–*LOC132999031-like* units are present in common dab and several related flatfish species. The diagram is not to scale for the lengths of each gene nor the distance between genes. Genomic information was retrieved from NCBI and Ensembl.

**S3 Fig. Coding-sequence alignment of the four intact common dab *gsdf* copies.** The alignment includes *gsdf1* and *gsdf2* on chromosome 1, *gsdf3* on chromosome 5 and the Y-linked *gsdfY* copy on chromosome 16. All four coding sequences are 600 bp long and contain no insertions, deletions or premature stop codons. Conserved positions are shown with a blue background, whereas variable positions are unshaded.

**S1 Table. Pairwise comparisons of the coding and predicted protein sequences of *gsdfY*, *gsdf1*, *gsdf2* and *gsdf3*.** Nucleotide and amino acid differences indicate the number of differing positions between the compared sequences. Identity values are expressed as percentages. Bold values indicate the highest CDS and protein identity with *gsdfY*.

## Notes

### Competing Interest Statement

The authors have declared no competing interest.

